# Structural and Energetic Details for the Formation of cGAS-DNA Oligomers

**DOI:** 10.1101/2023.09.03.556085

**Authors:** Xiaowen Wang, Wenjin Li

## Abstract

Upon binding to cytosolic DNA, the cyclic GMP-AMP synthase (cGAS) is activated to catalyze the synthesis of cGAMP, which then activates downstream effectors and induces innate immune responses. The activation of cGAS relies on the formation of cGAS-DNA oligomers and liquid phase condensation, which are sensitive to the length and concentration of DNA. For a thorough understanding of such a length-and concentration-dependent activation, the details of the cGAS-DNA oligomerization are required. Here, with molecular dynamics (MD) simulations, we report the structure of the cGAS-DNA monomer (the cGAS_1_-DNA_1_ complex), in which the DNA binds simultaneously to the major parts of two DNA-binding sites as observed in the cGAS-DNA dimer (the cGAS_2_-DNA_2_ complex) and the active site is largely immature. Energetic analysis reveals that two cGAS_1_-DNA_1_ complexes are just slightly less stable than the cGAS_2_-DNA_2_ complex and the energy barrier for the formation of cGAS_2_-DNA_2_ complex from two cGAS_1_-DNA_1_ complexes is high, suggesting that cGAS-DNA oligomerization is unfavored thermodynamically and kinetically in low concentration of cGAS and DNA. However, the formation of cGAS_4_-DNA_2_ complex from one molecule of cGAS_2_-DNA_2_ complex between cGAS and long DNA and two molecules of cGAS are energetically favored without energy barrier.

## Introduction

In the innate immune response, the body detects the presence of pathogenic nucleic acids to activate downstream immune responses and defend against invading pathogens. The host cell recognizes nucleic acids present in different subcellular compartments via specific sensors or pattern-recognition receptors.^1^ For example, toll-like receptors detect the nucleic acids in endosomes and retinoic acid-inducible gene I–like receptors recognize cytosolic RNAs. In 2013, the cyclic GMP–AMP synthase (cGAS) was identified as the major cytosolic DNA sensor.^2^ cGAS is a nucleotidyl transferase and is activated upon DNA binding to catalyze the synthesis of cyclic GMP–AMP (cGAMP), which then activates the adaptor protein stimulator of interferon genes (STING) in the endoplasmic reticulum. STING induces the expression of the type I interferons and other cytokines.^3,4^ Thus, the recognition of cytosol DNA by cGAS is one of the key steps to trigger innate immune responses.

The structures of cGAS-DNA complexes were then resolved soon by several research groups and provided the structural basis of cytosol DNA sensing by cGAS. ^5–8^ The activation of cGAS requires the formation of oligomers between cGAS and DNA. ^5^ The crystal structure of mouse cGAS (mcGAS) bound to an 18 bp DNA (PDB ID: 4LEY)^5^ showed that the activated cGAS forms a 2:2 complex with DNA (cGAS_2_-DNA_2_ complex) instead of the previously reported 1:1 complex (cGAS_1_-DNA_1_ complex). ^6,7^ In the cGAS_2_-DNA_2_ complex, each cGAS interacts with two DNA molecules via two binding sites, sites A and B, respectively (Fig. 1a). Compared to its inactive apo-state, ^6^ the activated cGAS by DNA-induced oligomerization undergoes significant conformational changes, which occur mainly in the spline helix at the DNA binding site A, the *β*5*/β*6*/β*7*/β*2 sheets where the catalytic residues locate, and the activation loop at the active site (Fig. 2a). ^5^ Although cGAS in the cGAS_1_-DNA_1_ complex is generally considered to be inactive, its structural basis remains unclear.

**Figure 1.**
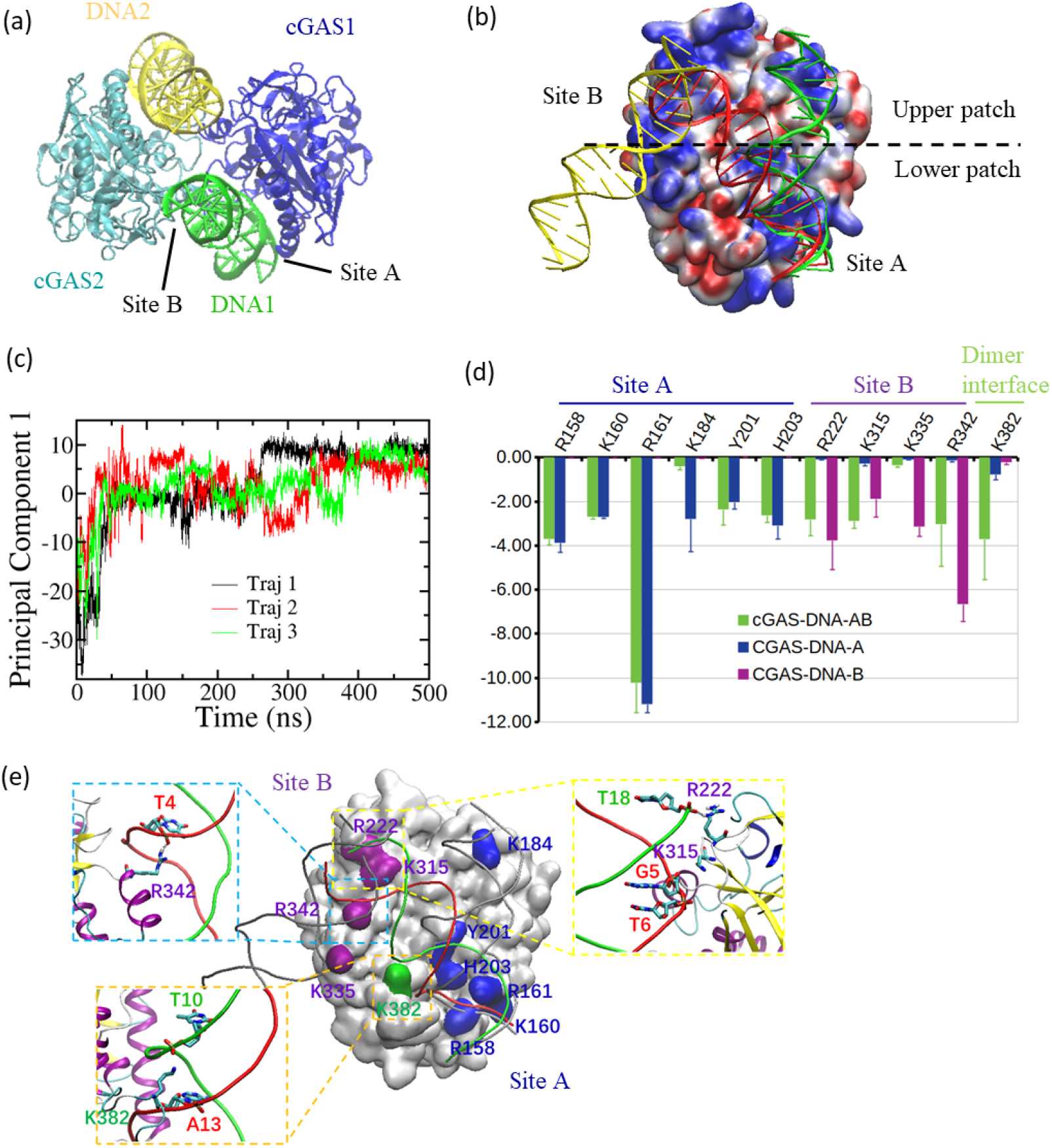
Structure and energetics of DNA in the cGAS_1_-DNA_1_ complex. (a) Crystal structure of 2:2 mcGAS:dsDNA complex (PDB ID: 4LEY^5^) is shown in a cartoon representation. cGAS1 (blue): chain A; cGAS2 (cyan); chain C; DNA1 (green): chains E and F; DNA2 (yellow): chains I and J. (b) Orientation of DNA (red cartoon) in the cGAS_1_-DNA_1_ complex when cGAS is fitted to cGAS1 in the crystal structure of the cGAS_2_-DNA_2_ dimer. cGAS1 is shown by a surface model colored with its electrostatic surface potential as a monomer. DNA1 and DNA2 bound to cGAS1 are shown with the same modes as the ones in (a). cGAS1, DNA1, and DNA2 are in their conformations in the dimer form. (c) Changes of the first principal component along with time in three parallel simulations of the cGAS_1_-DNA_1_ complex, starting from the conformation of cGAS1 and DNA1 in the crystal structure of the cGAS-DNA dimer. (d) Results of MM/GBSA analysis for important residues in 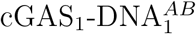 (green bars), cGAS with DNA bound to site A (blue bars), and cGAS with DNA bound to site B (purple bars). Error bars are the standard deviation over three simulations. (e) Locations of important residues listed in (d) and the interaction details for selected residues with DNA. An average structure of 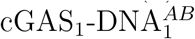 is shown with cGAS by surface mode and DNA by tube mode. The Waston strand of DNA is in red, while the Crick strand is in green. DNA1 and DNA2 in their conformations in the dimer form are shown in grey tubes. Residues located at site A, site B, and dimer interface are shown in blue, purple, and green, respectively.

**Figure 2.**
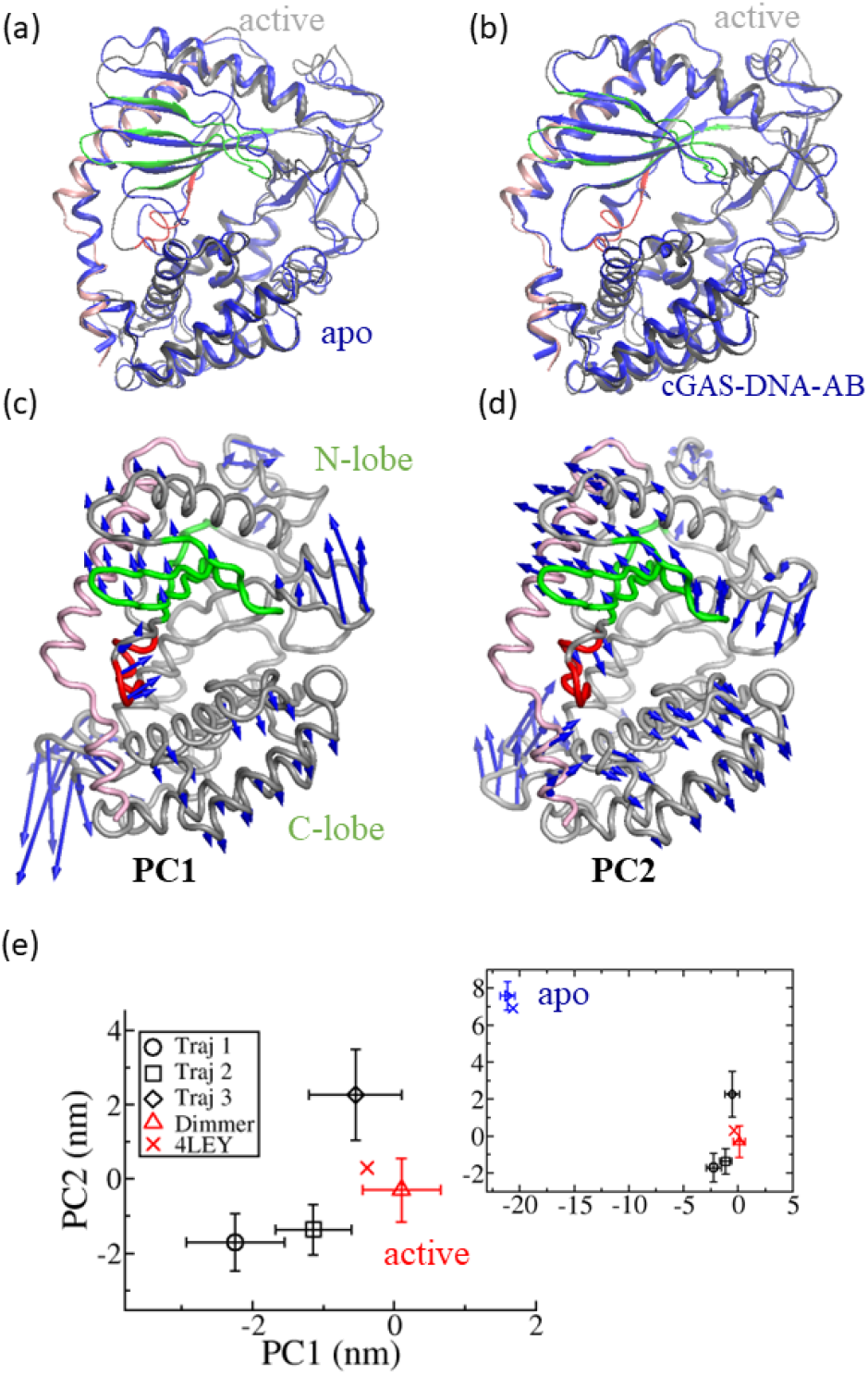
Structure of the active site of cGAS in various forms. (a) Structural comparison of the apo cGAS (blue, chain A in PDB ID: 4K8V^6^) and the active cGAS (grey, chain A in PDB ID: 4LEY) with the spine helix (*α*1*/α*2), the *β*5*/β*6*/β*7*/β*2 sheets, and the activation loop being highlighted in pink, green, and red, respectively. (b) Structural comparison of cGAS in 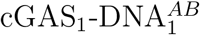 (blue) and the active cGAS (the same as the one in (a)). (c) and (d) visualize the motions of the first (PC1) and second (PC2) principal components from PCA on all the three simulations of the cGAS_1_-DNA_1_ complex. (e) Projection of cGAS structures onto the first two principal components. The average position of cGAS in 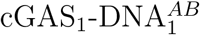 in each simulation is calculated from the last 200 ns of trajectories (black circle, square, and diamond); The average position of cGAS in the cGAS_2_-DNA_2_ dimer are evaluated over three parallel simulations (red triangle up); blue triangle right is the position of the apo cGAS averaged over a 500 ns simulation; blue and red cross correspond to cGAS in the crystal structures of apo form and cGAS-DNA dimer, respectively.

The activity of cGAS depends on the length of DNA and the ability of long DNA to activate cGAS is generally higher than the one of short DNA in both *in vitro* and *in vivo* assays.^9,10^ 20 bp DNA can activate cGAS through the formation of cGAS_2_-DNA_2_ complexes *in vitro*, it however is largely not immunostimulatory *in vivo*.^9,11^ The activation of cGAS *in vivo* required DNA of length no less than 45 bp. cGAS and DNA of 39 bp form a cGAS_4_-DNA_2_ complex, in which two cGAS dimers along DNA are ‘head-to-head’ oriented.^9^ The DNA length-dependent activation of cGAS can be explained by a ‘DNA-protein ladder’ model: the binding of cGAS along DNA was suggested to be in a stepwise fashion; the assembly of the first cGAS dimer with two DNA molecules into a cGAS_2_-DNA_2_ complex is not effective, however the cGAS_2_-DNA_2_ complex pre-align two DNA into ladder sides that facilitates the binding of subsequent cGAS dimers. ^9^ In addition, DNA binding to cGAS was shown to robustly induce phase transition to form liquidlike droplets in a concentration- and length-dependent manner.^12^ The formation of protein-DNA ladder could be a vital step for the formation of liquidlike droplets.^13^ However, how such cGAS-DNA oligomers assemble is not fully understood. The elucidation of the dynamical details of these assemblies will deepen the understanding of the mechanism by which DNA-dependent activation of cGAS is delicately regulated in a concentration-and length-dependent manner to ensure its proper function in innate immunity.

Here, molecular dynamics (MD) simulations^14^ were performed to characterize the formation of cGAS-DNA oligomers from their monomers with mcGAS as a typical example. In the following, we refer mcGAS with cGAS unless specifically stated. We observe that the DNA binds to the major patches of DNA-binding sites A and B simultaneously in the cGAS_1_-DNA_1_ complex, which is quite different to the monomers seen in the cGAS_2_-DNA_2_ complex. The cGAS is demonstrated by structural analysis to be largely not engaged in a fully active state, which provides the structural basis for the inactivity of the cGAS_1_-DNA_1_ complex. Energetic analyses shown that the conformation of the cGAS_1_-DNA_1_ complex is more stable than the one seen in the cGAS_2_-DNA_2_ complex and the cGAS_1_-DNA_1_ is suggested to be the main species in case of short DNA, consistent with previous experiments. ^9^ The assembly of the cGAS_2_-DNA_2_ complex from two cGAS_1_-DNA_1_ complexes was suggested to be most favorable, ruling out the possibility to form this complex from cGAS dimer and two parallel DNA molecules. Force-probe MD simulations^15,16^ reveal the disassembly details of the cGAS_2_-DNA_2_ complex in case of both short DNA and long DNA, which indicates that the prearrangement of long DNA upon the formation of the cGAS_2_-DNA_2_ complex facilitate the binding of subsequent cGAS. Collectively, these results provided structural and energetic details of how the cGAS-DNA oligomers form and rationalized the cooperative DNA sensing of cGAS in a concentration- and length-dependent manner.

## Results

### DNA binding interface in the cGAS_1_-DNA_1_ complex

It is reasonable to doubt that the cGAS_1_-DNA_1_ complex adopts the same conformation as the one in the cGAS_2_-DNA_2_ complex. We thus started from the crystal structure of cGAS_1_-DNA_1_ complex in the cGAS_2_-DNA_2_ complex and performed three parallel simulations of 500 ns. All these simulations end in a conformation, in which the DNA rotates about 40 degree relative to its conformation in the crystal structure (Fig. 1b). As a result, DNA binds to both sites A and B and the DNA binding interface consists of the lower patch of site A and the upper patch of site B. The conformation is considered to be the most stable structure for the cGAS_1_-DNA_1_ monomer and is named as 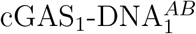. During all the simulations, one end of the DNA moves away from the spine helix and binds with the loop connecting the stands *β*2 and *β*3 (Movie S1), which can be characterized by the first principal component from a principle component analysis (PCA) on the trajectories of three simulations (Fig. S1 and S2A). The changes of the first principal component over time (Fig. 1c) show that the structures of the cGAS_1_-DNA_1_ monomer switch to 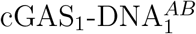 within 100 ns and remain there for the rest of the simulations, indicating that 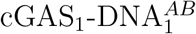 is a stable conformation. To identify the important interactions between cGAS and DNA in 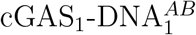, we performed MM/GBSA (molecular mechanics generalized-Born surface area) analysis for the last 300 ns of the three simulations.^17,18^ The energetic contributions of important residues in 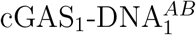 are shown in Fig. 1d (green bars). As a comparison to the interactions in the cGAS_2_-DNA_2_ dimer, the interactions between cGAS and DNA at the site A (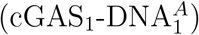) and the one between cGAS and DNA at site B (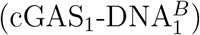) are also calculated from MM/GBSA analysis on the MD simulations of the cGAS_2_-DNA_2_ dimer (blue and purple bars in Fig. 1d). In the 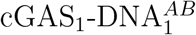, DNA interacts with R158, K160, R161, Y201, and H203 at the lower patch of site A with almost the same strength as the ones in the 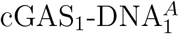 of the cGAS_2_-DNA_2_ dimer, while it loses the interaction with K184 at the upper patch of site A. DNA in 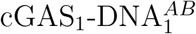 binds the upper path of site B with an orientation that is almost perpendicular to the one of DNA in 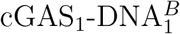 (Fig. 1b). At the site B, DNA interacts with R222, K315, and R342 at the upper patch. The interaction of DNA with R342 in 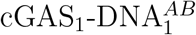 is about -3.0*±*1.9 kcal/mol, which is much lower in magnitude than the one of -6.7*±*0.8 kcal/mol in 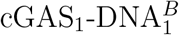. The interaction of DNA with R222 is significantly weaker than the one in 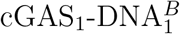, while the interaction of DNA with K315 is slightly stronger than the one in 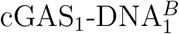. The DNA in 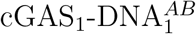 has no interaction with K335 at the lower patch of site B. In 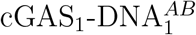, R342 binds from the major groove side with T4 in the Watson strand; R222 binds with T18 in the Crick strand and K315 interacts with G5 and T6 in the Watson strand from the minor groove side (Fig. 1e). Interestingly, DNA also binds strongly with K382 via T10 in the Crick strand and A13 in the Watson strand from the minor groove side (Fig. 1e), while DNA does not interact with K382 in the cGAS_2_-DNA_2_ dimer (Fig. 1d).

### Structure of the active site in the cGAS_1_-DNA_1_ complex

As DNA is suggested to form a 1:1 cGAS:DNA monomer with a binding interface that is very different to the one in the cGAS_2_-DNA_2_ dimer, where the active site of cGAS is in an active state, we thus wonder whether the active site undergoes significant conformational changes. By comparing the structure of cGAS in 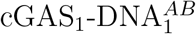 with the one (cGAS1) in the crystal structure of the cGAS_2_-DNA_2_ dimer (Fig. 2b), the conformation of the activation loop is observed to be significantly different to the one in the active state and forms a short helix, the *β*1 sheet becomes shorter with the amino acids next to the activation loop switches to loop structure. The *β*5*/β*6*/β*7*/β*2 sheets exhibits distinguishable shifts as compared with the conformation in the active state. To further unveiling the conformational differences between the cGAS in 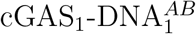 and the one in the active form, PCA was performed for cGAS only on the trajectories of three simulations of the cGAS_1_-DNA_1_ complex. The conformational changes of cGAS can be characterized by the first two principal components (Fig. S2B), which is shown in Figs. 2c and 2d. The first principal component describes the outward movement of the short helix in the activation loop and the upward shift of the *β*5*/β*6*/β*7*/β*2 sheets; The second principal component involves the change of amino acids in the activation loop connecting the *β*1 sheet and the movement of the *β*5*/β*6*/β*7*/β*2 sheets towards the left. The N-lobe and the C-lobe of cGAS move in an opposite direction in the two principal components. We then project the conformation of cGAS in the apo form, the conformation of cGAS in the active state and the ones in 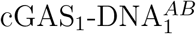 onto the two principal components (Fig. 2e). The conformations of cGAS in 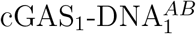 from the three independent simulations are observed to distinct from its active form, although they are all different to each other. Compared to the conformation in the apo form, the conformation of cGAS in 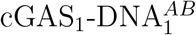 are very close to the one in the active state (inner plot of Fig. 2e). As the 1:1 cGAS:DNA monomer is largely suggested to be inactive, our results indicate that subtle changes in the active site of cGAS in 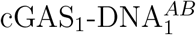 reduce the catalytic activity to a great extent.

### Energetics of various complexes in oligomerization from MM/GBSA analysis

To understand the mechanism of assembly of cGAS and DNA into oligomers, we calculated the free energies for the formation of various complexes in oligomerization with the MM/GBSA method, the relative free energies (RFEs) of these complexes compared to the free energy of free cGAS and/or DNA. We first examined the energetics for the formation of various binary complexes from cGAS and/or DNA. As shown in Table 1, the RFE for the formation of 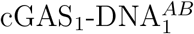 is about -146*±*21 kcal/mol, which is significantly higher in magnitude than the one for 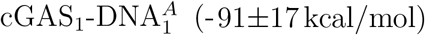. It suggests that cGAS and DNA prefer to forming the structure of 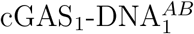 rather than 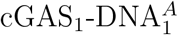, the conformation observed in the 2:2 cGAS:DNA complex. It also consistent with the results of MD simulations, where the structure of 1:1 cGAS:DNA complex switches from 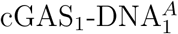 to 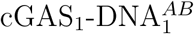 in all the three simulations. The RFE of 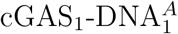 is lower than the one of 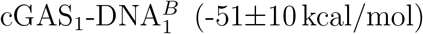, suggesting that the interaction between cGAS and DNA at the DNA binding site A is stronger than the one at the site B. The formation of cGAS:cGAS dimer is energetically favored with a RFE of -30*±*12 kcal/mol and residues K382, I380, and E381 at the dimer interface contribute most to the interaction with per-residue binding free energies of -2.62*±*0.25 kcal/mol, -1.86*±*0.21 kcal/mol, and -1.80*±*0.24 kcal/mol, respectively. The interaction between two DNAs in the cGAS_2_-DNA_2_ complex is weakly unfavored with a RFE of 15*±*2 kcal/mol. Among all the binary complexes, the RFE of 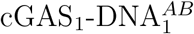 is the lowest, indicating that cGAS_1_-DNA^*AB*^ is the most favored binary complex.

**Table 1:** Relative free energies (RFEs) for the formation of various complexes in cGAS-DNA oligomerization. The results are obtained from MM/GBSA analysis on both the simulations of the cGAS-DNA dimer and the ones of the cGAS-DNA monomer with entropies being taken into account.

| index | Formation of complexes | RFE (kcal/mol) |
| --- | --- | --- |
| 1 | cGAS<sub>1</sub>-DNA<sub>1</sub><sup>AB</sup> | -146±21 |
| 2 | cGAS<sub>1</sub>-DNA<sub>1</sub><sup>A</sup> | -91±17 |
| 3 | cGAS<sub>1</sub>-DNA<sub>1</sub><sup>B</sup> | -51±10 |
| 4 | cGAS<sub>2</sub> | -30±12 |
| 5 | DNA<sub>2</sub> | 15±2 |
| 6 | cGAS<sub>1</sub>-DNA<sub>2</sub> | -139±20 |
| 7 | cGAS<sub>2</sub>-DNA<sub>1</sub> | -172±24 |
| 8 | cGAS<sub>2</sub>-DNA<sub>2</sub> | -313±31 |

The energetics for the formation of ternary and quaternary complexes are also analyzed. The RFE for the formation of the ternary complex cGAS_1_-DNA_2_ from one cGAS and two DNA molecules is -139*±*20 kcal/mol, and the one for the formation of the ternary complex cGAS_2_-DNA_1_ from two cGAS and one DNA molecules is -172*±*24 kcal/mol. In addition, the RFE for the formation of the cGAS_2_-DNA_2_ dimer from two cGAS and two DNA molecules is estimated to be -313*±*31 kcal/mol.

### Disassembly of the cGAS_2_-DNA_2_ complex

As suggested by the results from MM/GBSA analysis, the binding affinity of DNA to cGAS at the site A in the cGAS_2_-DNA_2_ complex is stronger than the one at the site B. Thus, the cGAS_2_-DNA_2_ complex is expected to disassemble first at the DNA binding site B. To validate this expectation, the early stage of disassemble of the cGAS_2_-DNA_2_ complex was simulated by force-probe MD simulations, in which the center of mass of each cGAS molecule was pulled under a constant rate. Two sets of simulations (three each) were carried out: one set without any restraints to mimic the disassembly of the cGAS_2_-DNA_2_ complex formed between cGAS and short DNA; one set with distance restraints to two DNAs to maintain the conformation of two DNAs in the cGAS_2_-DNA_2_ complex, which mimic the dissociation of cGAS from the cGAS_4_-DNA_2_ oligomer formed between cGAS and long DNA, where the two DNAs are aligned and thus restrained by the other cGAS_2_-DNA_2_ complex. In all the simulations, the dissociation of DNA from cGAS occurs at the DNA binding site B. These results confirmed that the binding of DNA to cGAS at the site A is stronger than the one at the site B. Note that there are two DNA binding sites B in the cGAS_2_-DNA_2_ complex and the cGAS-DNA interaction may rupture randomly at one of them. Interestingly, instead of dissociating at one of the B-sites, the cGAS and DNA in the set of simulations without any restraints loses part of their interactions at both B-sites, where the interaction between DNA and helix *α*6 breaks (Fig. 3a, movie S2). In this intermediate (named INT1), the C-lobes of two cGAS molecules move far away from each other and the N-lobes come close. As a result, the two DNA molecules are almost perpendicular to each other. In the set of simulations with the conformation of two DNAs being restrained, one of the two cGAS molecules in the cGAS_2_-DNA_2_ complex separates completely from DNA at the B-site, while its interaction with DNA at the A-site is large intact (named INT2, Fig. 3b, movie S3). The intermediates INT1 and INT2 during the disassembly of the cGAS-DNA dimer in case of free and restrained DNAs are quite different, suggesting that the disassembly pathway of the cGAS_2_-DNA_2_ complex for short DNAs could be distinct from the one of the cGAS_4_-DNA_2_ oligomer for long DNAs. MM/GBSA analyses of the two intermediates show that the RFE of INT1 is -260*±*23 kcal/mol and the RFE of INT2 is -229*±*25 kcal/mol, which is about 31 kcal/mol higher than the one of INT1. The lower RFE of the INT1 than the INT2 suggests that the dimer between cGAS and short DNA breaks via a pathway of much lower energy barrier than the one in the cGAS_4_-DNA_2_ oligomer.

**Figure 3.**
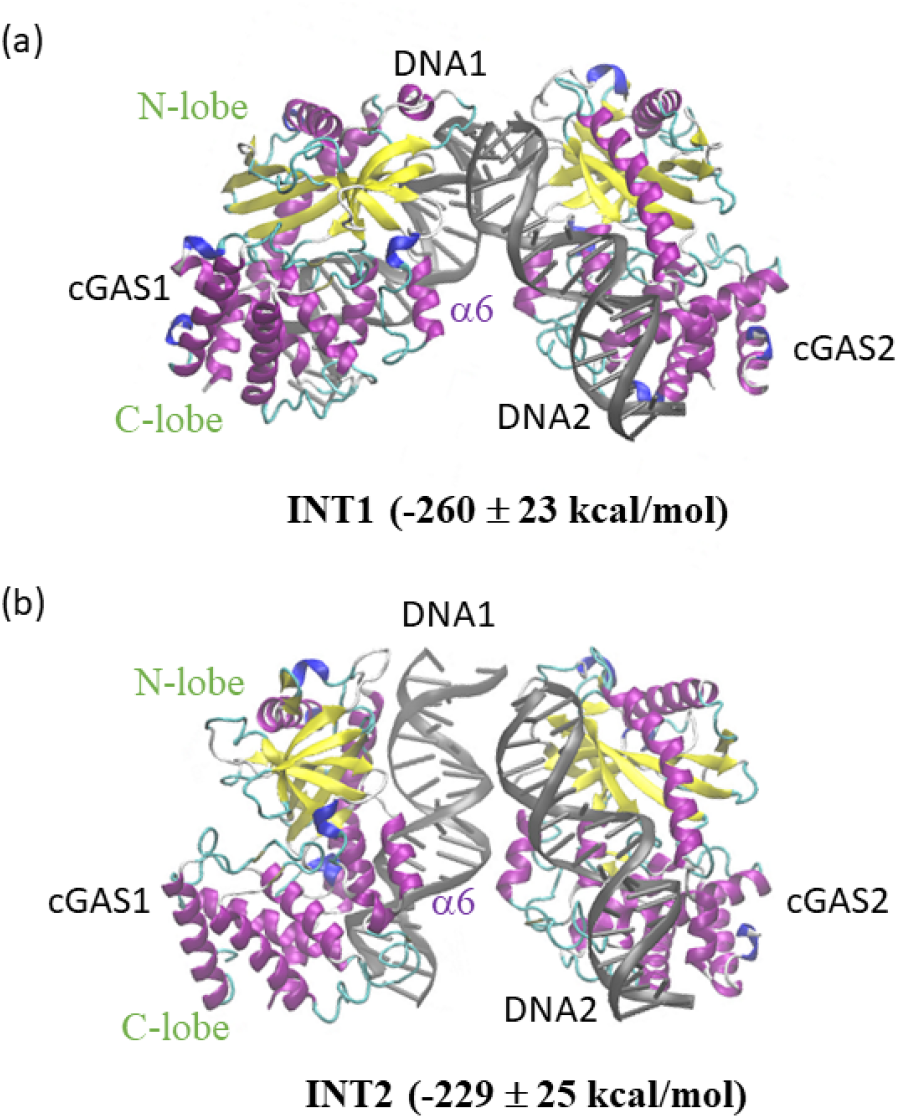
Intermediates in the force-induced disassembly of the cGAS-DNA dimer from two sets of molecular dynamics simulations: (a) a set without any restraint; (b) a set with distance restraints to two DNAs. cGASs are colored according to their secondary structures and DNAs are in grey.

## Discussions

The activation of cGAS by DNA requires the formation of a cGAS-DNA oligomer and the cGAS-DNA monomer is not active,^5,9,19^ but the structural basis remains not fully understood. Here, we proposed a structural model for the cGAS_1_-DNA_1_ complex, in which the DNA binding interface is quite different to the one in the cGAS-DNA oligomer and the conformation of active site of cGAS deviates from the one in an active form. Thus, insufficiency of the cGAS-DNA monomer to active cGAS as a nucleotidyl transferase is supported with details at the atomic level. 11 residues (R158, K160, R161, K184, Y201, H203, R222, K315, K335, R342 and K382) are suggested to be important for the activation of cGAS (Fig. 1d), which is largely in agreement with previous experimental and computational studies. ^5,20^ These residues may be divided into three categories: the first category that is essential for the formation of the cGAS-DNA dimer, the second one that is required to hold the conformation of the active site in the right position, and the third one that fall into both the first and the second categories. The first category includes residues R158, K160, R161, K184, K335, and R342, which interact strongly with DNA in the cGAS_2_-DNA_2_ complex. K382 is a key residue that mediates cGAS-cGAS interaction and belongs to the first category as well. Residues Y201 and K315 belong to the second category, while residues R222 and H203 fall in the third one. R222 locates at the loop connecting strands *β*2 and *β*3, K315 sits in the loop between the strand *β*7 and helix *α*5. The conformation shifting of the *β*5*/β*6*/β*7*/β*2 sheets in the switch from 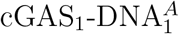 to 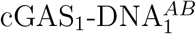 may be caused by the changes in the interaction of the two residues with DNA. Note that, the DNA bound to these residues in 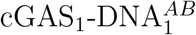 with an orientation that differs by about 90° to the one in 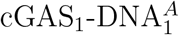. Residues Y201 and H203 are located in the activation loop and their positional changes with the rotation of the DNA in the transition from 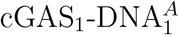 to 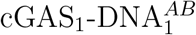 induce the conformational change of the activation loop. These conformational changes in the *β*5*/β*6*/β*7*/β*2 sheets and the activation loop are suggested to render together the cGAS in cGAS_1_-DNA_1_ complex to be inactive to a great extent. Since the DNA binds in part to both the sites A and B to form the 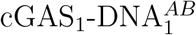 complex, it excludes the bind of other DNA to cGAS and thus the formation of the cGAS_2_-DNA_2_ complex. In addition, DNA binds strongly with R382 in the 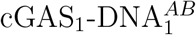 complex, which hinders the dimerization of the cGAS-DNA binary complexes as well.

The positive electrostatic surface potential (ESP) of cGAS at the dimer interface between two cGAS-DNA monomers distributes mainly at the DNA binding sites A and B, which is separated by negative ESP in between (Fig. 1b). These regions of positive ESP can be further divided into four patches: the lower and upper patches of the A-site, and the lower and upper patches of the B-site. The lower patch of the A-site and the upper patch of the B-site are two major patches for protein-DNA interaction, as five important residues (R158, K160, R161, H201, and H203) locate in the lower patch of the A-site and three important ones (R222, K315, and R342) sit in the upper patch of the B-site (Fig. 1e). The upper patch of the A-site and the lower patch of the B-site are two minor patches, with one important residue locating on each of them. Residues R222 and K315 in one of the major patches (the upper patch of the B-site) connect closely with the *β*5*/β*6*/β*7*/β*2 sheets where the catalytic residues locate, while Residues Y201 and H203 in the other major patch sit right in the activation loop. These residues setup the connection between the binding of DNA to the major patches and conformational changes of the active site. Such a brilliant design of the ESP enables the fine-tuning of cGAS activity by DNA and could be a result of natural selection.

With the knowledge of the energetics of various complexes in oligomerization (Table 1), the thermodynamic and kinetic aspects for the protein-DNA oligomerization could be deduced (Fig. 4). Consistent with the DNA-protein ladder model proposed in a previous study, ^9^ 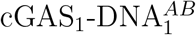 is suggested to be formed first in the cGAS-DNA oligomerization, as its formation is most favored among all the possible binary complexes. Then the DNA moves away from the upper patch of the B-site to form the 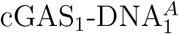, which in turn assembles into the cGAS_2_-DNA_2_ complex (Fig. 4a). The RFE of the two 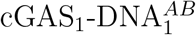 complexes (-292*±*30 kcal/mol) is slightly higher than the one of cGAS_2_-DNA_2_ complex (-313*±*31 kcal/mol). Thus, the cGAS-DNA monomer is highly populated in the presence of short DNA. However, under high concentration of cGAS and short DNA, the concentration of the cGAS_2_-DNA_2_ complex can be high enough and thus cGAS can be activated by short DNA *in vitro*. In the case of long DNA, the cGAS_2_-DNA_2_ complex can be further stabilized with the subsequent binding of cGAS to the parallelized DNA and thus cGAS can be activated by long DNA at low amounts. From a kinetical viewpoint, the cGAS_2_-DNA_2_ complex is unstable as the energy barrier for its disassembly into cGAS-DNA monomers is relatively low (∼131 kcal/mol). As a comparison, the dissociation of one cGAS from the cGAS_4_-DNA_2_ complex requires to get over a much higher energy barrier of 174 kcal/mol. Thus, the life-time of active cGAS in the cGAS_4_-DNA_2_ complex is considerably longer than the one in the cGAS_2_-DNA_2_ complex. Furthermore, short DNA can form active cGAS_2_-DNA_2_ complex, however it may be trapped in the inactive 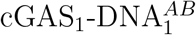 state kinetically due to the high activation barrier. cGAS can quickly binds parallelized long DNA to form active protein-DNA ladder with no energy barrier, where the parallelized DNA can be provided by the cGAS_2_-DNA_2_ complex or cellular factors such as HMGB proteins. ^10^ These results could be the explanation for the fact that long DNA instead of short DNA is able to activate the cGAS pathway *in vivo*.^9–11^

**Figure 4.**
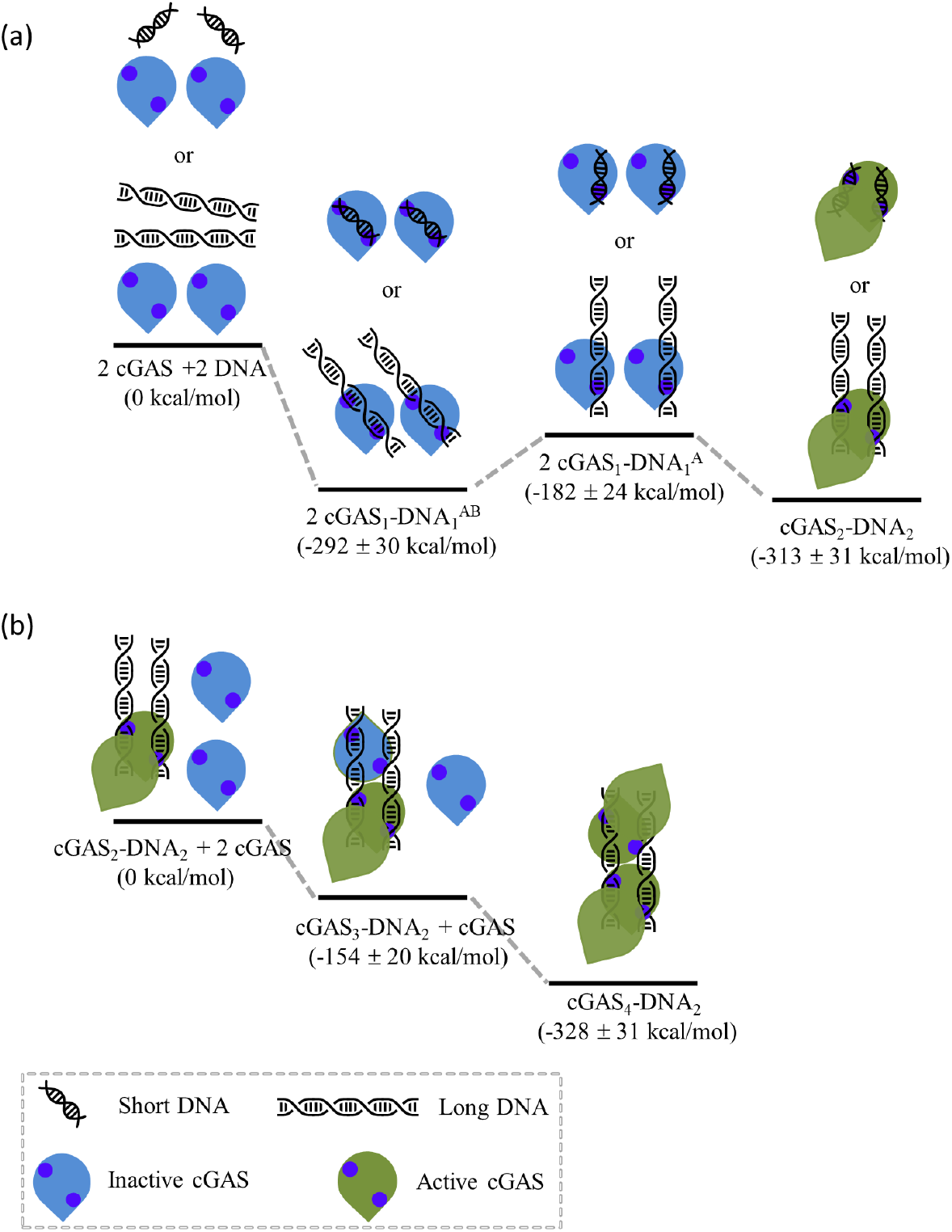
Assembly scheme for the cGAS-DNA oligomer. (a) Short or long DNA binds to cGAS to form 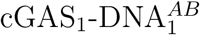, these 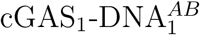 complexes then assemble into the cGAS-DNA dimer via a transition state of the 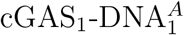 structures. (b) In the case of long DNAs, two cGAS molecules bind sequentially to the parallelized DNA in the cGAS_2_-DNA_2_ complex to form the cGAS_4_-DNA_2_ complex. Here, the complexes cGAS_2_-DNA_2_, cGAS_3_-DNA_2_, and cGAS_3_-DNA_2_ are analogue to DNA_2_, cGAS_1_-DNA_2_, and cGAS_2_-DNA_2_ in table 1, from which the RFEs of these complexes are derived. The first states in (a) and (b) are used as the reference states. Two blue circles represent the upper patch of B-site and the lower patch of A-site.

Mutations N172K and R180L of mcGAS reduced the amount of cGAMP production in the presence of short or long DNA.^10^ These mutations may lower interactions of the DNA with the upper patch of A-site and increase the RFE of the cGAS_2_-DNA_2_ and 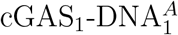 complexes, while the one of 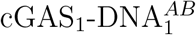 is unaffected. Consequently, the two substitutions reduce the stability of the cGAS-DNA dimer and raise the energy barrier for the formation of the dimer from two 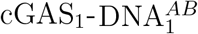 complexes. Thus, the formation of the cGAS_2_-DNA_2_ complex between this mutant of mcGAS with short DNA is thermodynamically and kinetically less favorable compared to wild type mcGAS. This may explain the enhanced DNA-length specificity of human cGAS compared to mcGAS as well. In the presence of short DNA, the molar ratio of DNA:cGAS was found to be 1.28 in an isothermal titration calorimetry of mcGAS and DNA,^9^ which is higher than the molar ratio of 1 in the 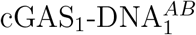 or cGAS_2_-DNA_2_ complexes. A plausible explanation could be that the cGAS_1_-DNA_2_ complex (DNA:cGAS molar ratio of 2) can also form to a great amount in the presence of excess short DNA, as the RFE of the cGAS_1_-DNA_2_ complex is marginally higher than the one of the 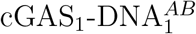 complex (Table 1).

In the disassembly of the cGAS_2_-DNA_2_ complex, the protein-DNA interactions break exclusively at the DNA binding B-site, at which the protein-DNA interaction is much weaker than the one at the A-site. One would expect a sequential breaking of the protein-DNA interactions at two B-sites in the cGAS_2_-DNA_2_ complex. Here, we observed that the two B-sites break partially to form an intermediate INT1 (Fig. 3a) in the force-probe MD simulations, indicating that the protein-DNA interaction at the lower patch of the B-site is weaker than the one at the upper patch. In the case of long DNA, the formation of the intermediate INT1 is less favored to the case of short DNA, as the two DNA molecules will clash with cGAS. Thus, the breaking of a cGAS_2_-DNA_2_ complex with long DNA is suggested to follow the sequential breaking of two B-sites. Thus, the disassembly pathway of the cGAS_2_-DNA_2_ complex could be length-dependent.

## Concluding Remarks

From extensive MD simulations and MM/GBSA analysis, we here proposed the structure of the cGAS-DNA monomer, which differ remarkably to its conformation in the cGAS-DNA dimer. The free energy of two cGAS-DNA monomers was found to be comparable to the one of the cGAS-DNA dimer. These results supported a previously proposed DNA-protein ladder model^9^ with structural and energetic evidences. The length- and concentration-dependent activation of cGAS by DNA could be due to the brilliant design of the ESP in the dimer interface, which consists of two major patches and two minor ones for protein-DNA interactions. Such a design results in various forms of protein-DNA interaction modes with these four patches, which corresponds to the diverse complexes in the cGAS-DNA oligomerization. Thus, the thermodynamics and kinetics for the processes in the cGAS-DNA oligomerization are fine-tuned by the length and concentration of DNA. Importantly, the conformation of the active site of cGAS is tightly coupled to the two major patches and the activity of cGAS is thus delicately controlled by DNA in a length- and concentration-dependent manner. The proposed results may shed light on the mechanism of other nucleic-acid-sensing pathways^21^ and facilitate the design of cGAS variants of therapeutical importance. ^22^

## Method

### Molecular Dynamics Simulation

All simulations were performed using the 2019 version of GROMACS program. ^23^ The AMBER14SB force field^24^ was used for cGAS and the parmbsc1 parameters^25^ for DNA. The zinc AMBER force field^26^ was used for the four-coordinated zinc metal center in the zinc-thumb domain and the TIP3P water model^27^ was used. The crystal structure (PDB ID: 4LEY)^5^ was used to as the starting model for the simulation of the cGAS-DNA dimer and its chains A, E, and F was selected to be the starting structure for the simulation of the cGAS-DNA dimer. The starting structure of apo cGAS was taken from the crystal structure (chain A in PDB ID: 4K8V). The protonation states of all titratable residues were carefully determined as in a previous study.^20^ The initial structures were hydrated in a cubic box of explicit waters and ions at the concentration of 100 mM. For each model, energy minimization with the maximum 5000 steps was carried out, followed by 100 ps MD simulation in the NVT ensemble and 200 ps simulation in the NPT ensemble with the position restraints on all heavy atoms. Then, with the final structure in the last simulation as the starting structure, 500 ns production runs were performed in the NPT ensemble. The systems were kept at constant temperature of 300 K and constant pressure of 1 bar with the velocity-rescaling thermostat^28^ and the Berendsen barostat,^29^ respectively. LINCS algorithm was used to constrain bonds involving hydrogen atoms^30^ and the integration time step was 2 fs. The force-switch modifier^31^ and particle mesh Ewald method^32^ were employed with a cut-off of 12 Å for van der Waals and long-range electrostatic interactions, respectively.

Starting with the final structures of production runs of the cGAS-DNA dimer, force-probe MD simulations^15,16^ were performed by pulling the center of mass of each cGAS molecule with a velocity of 0.003 nm/ns and a simulation time of 400 ns. In one set of the simulations, the conformation of two DNA was restrained with distance restraints on four distances, the distances of two ends of one DNA to the ones of the other DNA, to their initial values and a spring constant of 3000 kJ/(mol·nm^2^). The Molecular structures were visualized with VMD ^33^ and PyMOL.^34^

### MM/GBSA Analysis

The binding free energy between molecules in the simulated systems was evaluated with the MM/GBSA method^35^ and the GB-neck2 model with an internal dielectric constant of 10 was employed for its good performance in nucleic acid–protein complexes. ^36^ The MM/GBSA calculations were carried out with the gmx MMPBSA tool^37,38^ for structures every 1 ns. The last 400 ns trajectories for the apo cGAS and the cGAS-DNA dimer were analyzed, while the last 300 ns trajectories for the cGAS-DNA monomer and the last 50 ns for force-probe MD simulations of the cGAS-DNA dimer were used for MM/GBSA analysis. For each system, entropy was estimated from the normal-mode analysis of harmonic frequencies from 30 frames after energy minimization. ^39^

## Supplementary Material

See supplementary material for the supplemental Figures S1-S2, movies for an MD simulation of the cGAS-DNA monomer (Movie S1), force-probe MD simulations of the cGAS-DNA dimer with no restraints (Movie S2) and with restraints on DNA (Movie S3).

## Acknowledgments

This work was supported by Natural Science Foundation of Guangdong Province, China (Grant No. 2023A1515010471) and National Natural Science Foundation of China (Grant No. 31770777).

## References

[1] Yin, Q.; Fu, T.-M.; Li, J.; Wu, H. Annual review of immunology 2015, 33, 393–416.

[2] Sun, L.; Wu, J.; Du, F.; Chen, X.; Chen, Z. J. Science 2013, 339, 786–791.

[3] Wu, J.; Chen, Z. J. Annual review of immunology 2014, 32, 461–488.

[4] Chen, Q.; Sun, L.; Chen, Z. J. Nature immunology 2016, 17, 1142–1149.

[5] Li, X.; Shu, C.; Yi, G.; Chaton, C. T.; Shelton, C. L.; Diao, J.; Zuo, X.; Kao, C. C.; Herr, A. B.; Li, P. Immunity 2013, 39, 1019–1031.

[6] Gao, P.; Ascano, M.; Wu, Y.; Barchet, W.; Gaffney, B. L.; Zillinger, T.; Serganov, A. A.; Liu, Y.; Jones, R. A.; Hartmann, G.; others Cell 2013, 153, 1094–1107.

[7] Civril, F.; Deimling, T.; de Oliveira Mann, C. C.; Ablasser, A.; Moldt, M.; Witte, G.; Hornung, V.; Hopfner, K.-P. Nature 2013, 498, 332–337.

[8] Kranzusch, P. J.; Lee, A. S.-Y.; Berger, J. M.; Doudna, J. A. Cell reports 2013, 3, 1362–1368.

[9] Andreeva, L.; Hiller, B.; Kostrewa, D.; Lässig, C.; de Oliveira Mann, C. C.; Jan Drexler, D.; Maiser, A.; Gaidt, M.; Leonhardt, H.; Hornung, V.; others Nature 2017, 549, 394–398.

[10] Zhou, W.; Whiteley, A. T.; de Oliveira Mann, C. C.; Morehouse, B. R.; Nowak, R. P.; Fischer, E. S.; Gray, N. S.; Mekalanos, J. J.; Kranzusch, P. J. Cell 2018, 174, 300–311.

[11] Herzner, A.-M.; Hagmann, C. A.; Goldeck, M.; Wolter, S.; Kübler, K.; Wittmann, S.; Gramberg, T.; Andreeva, L.; Hopfner, K.-P.; Mertens, C.; others Nature immunology 2015, 16, 1025–1033.

[12] Du, M.; Chen, Z. J. Science 2018, 361, 704–709.

[13] Hopfner, K.-P.; Hornung, V. Nature reviews Molecular cell biology 2020, 21, 501–521.

[14] Karplus, M.; McCammon, J. A. Nature structural biology 2002, 9, 646–652.

[15] Grubmüller, H.; Heymann, B.; Tavan, P. Science 1996, 271, 997–999.

[16] Grubmüller, H. Protein-Ligand Interactions: Methods and Applications; Springer, 2005; pp 493–515.

[17] Hou, T.; Wang, J.; Li, Y.; Wang, W. Journal of chemical information and modeling 2011, 51, 69–82.

[18] Wang, E.; Sun, H.; Wang, J.; Wang, Z.; Liu, H.; Zhang, J. Z.; Hou, T. Chemical reviews 2019,119, 9478–9508.

[19] Zhang, X.; Wu, J.; Du, F.; Xu, H.; Sun, L.; Chen, Z.; Brautigam, C. A.; Zhang, X.; Chen, Z. J. Cell reports 2014, 6, 421–430.

[20] Wang, X.; Zhang, H.; Li, W. Physical Chemistry Chemical Physics 2020, 22, 26390–26401.

[21] Sohn, J.; Hur, S. Current opinion in structural biology 2016, 37, 134–144.

[22] Dowling, Q. M.; Volkman, H. E.; Gray, E. E.; Ovchinnikov, S.; Cambier, S.; Bera, A. K.; Sankaran, B.; Johnson, M. R.; Bick, M. J.; Kang, A.; others Nature Structural & Molecular Biology 2023, 30, 72–80.

[23] Abraham, M. J.; Murtola, T.; Schulz, R.; Páll, S.; Smith, J. C.; Hess, B.; Lindahl, E. SoftwareX 2015, 1, 19–25.

[24] Maier, J. A.; Martinez, C.; Kasavajhala, K.; Wickstrom, L.; Hauser, K. E.; Simmerling, C. Journal of chemical theory and computation 2015, 11, 3696–3713.

[25] Ivani, I.; Dans, P. D.; Noy, A.; Pérez, A.; Faustino, I.; Hospital, A.; Walther, J.; Andrio, P.; Goñi, R.; Balaceanu, A.; others Nature methods 2016, 13, 55–58.

[26] Peters, M. B.; Yang, Y.; Wang, B.; Fusti-Molnar, L.; Weaver, M. N.; Merz Jr, K. M. Journal of chemical theory and computation 2010, 6, 2935–2947.

[27] Jorgensen, W. L.; Chandrasekhar, J.; Madura, J. D.; Impey, R. W.; Klein, M. L. The Journal of chemical physics 1983, 79, 926–935.

[28] Bussi, G.; Donadio, D.; Parrinello, M. The Journal of chemical physics 2007, 126.

[29] Berendsen, H. J.; Postma, J. v.; Van Gunsteren, W. F.; DiNola, A.; Haak, J. R. The Journal of chemical physics 1984, 81, 3684–3690.

[30] Hess, B.; Bekker, H.; Berendsen, H. J.; Fraaije, J. G. J. Comput. Chem. 1997, 18, 1463–1472.

[31] Steinbach, P. J.; Brooks, B. R. J. Comput. Chem. 1994, 15, 667–683.

[32] Essmann, U.; Perera, L.; Berkowitz, M. L.; Darden, T.; Lee, H.; Pedersen, L. G. J. Chem. Phys. 1995, 103, 8577–8593.

[33] Humphrey, W.; Dalke, A.; Schulten, K. J. Mol. Graph. 1996, 14, 33–38.

[34] DeLano, W. L.; others CCP4 Newsl. Protein Crystallogr 2002, 40, 82–92.

[35] Tsui, V.; Case, D. A. Journal of the American Chemical Society 2000, 122, 2489–2498.

[36] Nguyen, H.; Perez, A.; Bermeo, S.; Simmerling, C. Journal of chemical theory and computation 2015, 11, 3714–3728.

[37] Valdés-Tresanco, M. S.; Valdés-Tresanco, M. E.; Valiente, P. A.; Moreno, E. Journal of chemical theory and computation 2021, 17, 6281–6291.

[38] Miller III, B. R.; McGee Jr, T. D.; Swails, J. M.; Homeyer, N.; Gohlke, H.; Roitberg, A. E. Journal of chemical theory and computation 2012, 8, 3314–3321.

[39] Genheden, S.; Kuhn, O.; Mikulskis, P.; Hoffmann, D.; Ryde, U. Journal of chemical information and modeling 2012, 52, 2079–2088.

