## supplemental Figures S1-S2 for "Structural and Energetic Details for the Formation of cGAS-DNA Oligomers"

### Supplementary Material

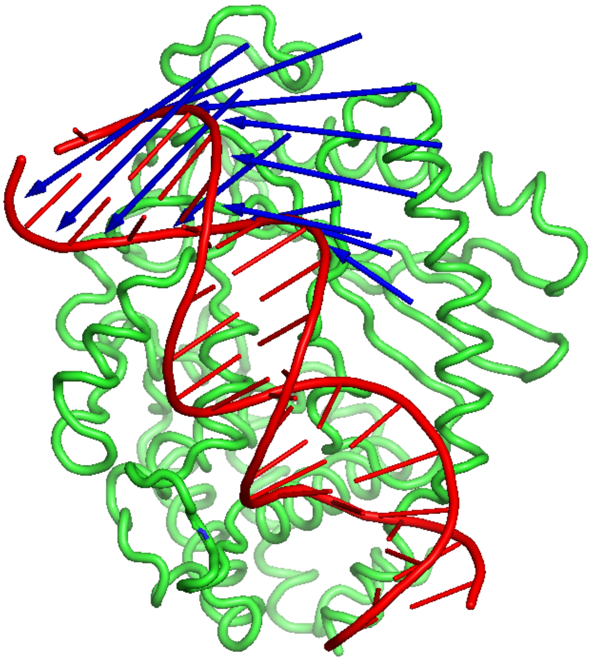

Figure S1: Visualize the motion of the first principal component from the principal component analysis of the cGAS-DNA monomer on all the three simulations of the $cGAS_1$-$DNA_1$ complex. cGAS is shown in green cartoon and DNA is shown in red cartoon. The movement of DNA is shown by blue arrows.

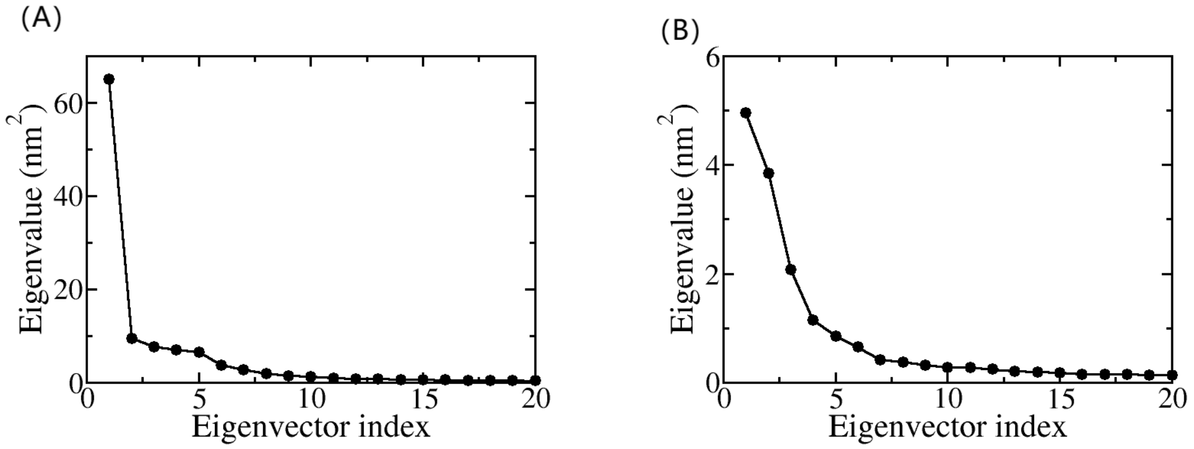


Figure S2: The eigenvalues of the first 20 eigenvectors for the principal component analysis of the cGAS-DNA monomer (A) or the cGAS only (B) on all the three simulations of the $cGAS_1$-$DNA_1$ complex.
